# The Effect of Bottleneck Size on Evolution in Nested Darwinian Populations

**DOI:** 10.1101/2022.09.22.508977

**Authors:** Matthew C. Nitschke, Andrew J. Black, Pierrick Bourrat, Paul B. Rainey

## Abstract

Previous work has shown how a minimal ecological structure consisting of patchily distributed resources and recurrent dispersal between patches can scaffold Darwinian properties onto collections of cells. When the timescale of dispersal is long compared with the time to consume resources, patches evolve such that their size increases, but at the expense of cells whose growth rate decreases within patches. This creates the conditions that initiate evolutionary transitions in individuality. A key assumption of this scaffolding is that a bottleneck is created during dispersal, so patches are founded by single cells. The bottleneck decreases competition within patches and hence creates a strong hereditary link at the level of patches. Here we construct a fully stochastic model of nested Darwinian populations and investigate how larger bottlenecks affect the evolutionary dynamics at both cell and collective levels. It is shown that, up to a point, larger bottlenecks simply slow the dynamics, but at some point, which depends on the parameters of the within-patch model, the direction of evolution toward the equilibrium is reversed. Introducing random bottleneck sizes with some positive probability of smaller sizes can counteract this, even if the probability of smaller bottlenecks is small.

## 1 Introduction

The biological world is a hierarchy of nested Darwinian populations, constructed through a series of major evolutionary transitions in individuality (ETIs) (Maynard Smith and Szathmáry, 1995; Calcott and Sterelny, 2012; Bourrat, 2019). How processes at one level affect others, both higher and lower, and the level at which selection acts are questions that have long occupied biologists (Lewontin, 1970; Damuth and Heisler, 1988; Sober and Wilson, 1998; Keller, 1999; Michod, 1999; Griesemer, 2000; Rainey et al., 2017; Herron et al., 2022) and philosophers (Okasha, 2008; Bouchard and Huneman, 2013; Clarke, 2014; Bourrat, 2021) alike. The evolution of new levels in the hierarchy poses a particular problem, as a mechanistic model must explain the emergence of Darwinian properties themselves, and not simply assume their existence (Griesemer, 2000; Okasha, 2008; Rainey and Kerr, 2010; Rainey et al., 2017). Recent work has drawn attention to the possibility that particular ecological conditions can exogenously impose Darwinian properties on collectives, thus causing higher levels, for example, collectives of cells, to participate in the process of evolution by natural selection in their own right. The idea, referred to as ecological scaffolding, is supported by both experimental (Hammerschmidt et al., 2014; Rose et al., 2020) and theoretical (Black et al., 2020; Doulcier et al., 2020) studies.

A simple model illustrates the idea of ecological scaffolding. Consider a population of cells where the resources needed for reproduction are divided into discrete patches. Multiple patches ensure patch-level discreteness and variation. Cell reproduction consumes resources and so periodic dispersal is required for long-term persistence. The dispersal process involves passage through a restrictive bottleneck with newly established patches being the offspring of parental patches. Cells are Darwinian by their inherent properties and manifest heritable variation in fitness (Lewontin, 1970; Godfrey-Smith, 2009). However, by virtue of ecological conditions (patchily distributed resources and a means of dispersal), patches are also Darwinian: patches vary one to another, patches reproduce (via dispersal) and offspring patches resemble parental patches (Black et al., 2020). Host-pathogen systems are another canonical example of ecologically scaffolded populations, with hosts corresponding to patches and transmission leading to dispersal and colonisation of new hosts being akin to a patch-level reproduction event (Gilchrist et al., 2002; André and Gandon, 2006; Coombs et al., 2007; Lythgoe et al., 2013).

When the period between dispersal events is long compared with the time for cells to consume resources, the composition of patches evolves such that patch fitness increases (larger patches at the time of dispersal are more likely to be the source of dispersing cells), but this leads to an apparent paradox: over the short term, cells that grow more slowly than the founding cells have an advantage because slower growing cells consume resources less rapidly. This tradeoff between improving patch (group) fitness and decreasing cell growth rate has been interpreted through the lens of fitness decoupling (Michod and Roze, 1999; Michod and Nedelcu, 2003; Okasha, 2005, 2008; Rainey and De Monte, 2014; Hammerschmidt et al., 2014), alluding to the fact that after an ETI, the fitness of the higher level construct (the patch) is no longer a simple function of the fitness of the individual components (the cells). However, the underlying assumptions of the notion of fitness decoupling and related terms such as “fitness transfer” or “export of fitness” have been called into question (Doulcier et al., 2022). In particular, it has been shown that properly measured, that is, measured over the same set of events, the fitness of cells and patches are always equal (Shelton and Michod, 2014; Bourrat, 2015b,a; Black et al., 2020). It has also been shown that, with the addition of a few assumptions, collectives under this framework can become resistant to the scaffolding being removed, thus resulting in a genuine ETI (Bourrat, in press). Moving on from the fitness decoupling concept, the dynamics observed during an evolutionary transition in individuality (ETI) have been interpreted in the context of a more general model involving tradeoff-breaking events (Bourrat et al., 2022).

A key to the model described so far is the bottleneck created by the dispersal process. Bottlenecks are a well-studied and important aspect of many developmental and evolutionary processes (Geoghegan et al., 2016; Grosberg and Strathmann, 2007; Melbinger et al., 2015; Kariuki et al., 2017; McCrone and Lauring, 2018; Nei et al., 1975). For example, there is a bottleneck created via the transmission of pathogens between hosts, and a bottleneck during multicellular reproduction. In the model described by Black et al (Black et al., 2020), patches are founded by single cells, so competition within patches is reduced (as all cells are related) and hence the composition of a patch is similar to the parent patch from which the colonising cell is dispersed. This creates a high degree of heritability (high correlation between parent and offspring phenotype) at the level of patches and hence facilitates a strong evolutionary response to selection at the higher level. This naturally generates questions as to the sensitivity of the ensuing evolutionary dynamics to bottleneck size. Providing answers stands to shed light on the importance of restrictive bottlenecks at the time of group-level reproduction and subsequent impacts of ETIs.

In this paper we construct a stochastic model of nested Darwinian populations and use this to explore how the number of cells dispersed affects the evolutionary dynamics of both cell and patch populations. Our model has the advantage of being mechanistic so the causes of different macroscopic dynamics can be transparently related back to the constituent parts of the system and their interactions. We concentrate our investigation on the regime where the length of time between dispersal events is long where increased patch size is generated at the expense of cell growth rate, creating a tension in the levels of selection.

We show that for bottlenecks bigger than one but still small, the evolutionary process that is induced by our ecology is slowed in its approach to the equilibrium, but is otherwise similar to a strict single cell bottleneck. After a point, the effect of increased competition within patches founded by multiple cells can overwhelm selection generated by the dispersal process at the level of patches and the direction of evolution from faster to slower growth rates is reversed. When the size of the bottleneck is changed from being fixed to a random variable with a distribution over possible sizes, we see lower level selection curtailed to some extent and the evolutionary equilibrium restored as long as there is some probability of smaller sizes.

## 2 Model

Figure 1 shows an overview of the model. The model consists of a fixed population of *M* patches, where each patch is initially seeded with some (small) number of cells. Cells replicate and mutate independently within each patch, but limited resources for growth, and the build-up of waste products, eventually leads to cell death becoming dominant and population size thus declines. Long-term persistence therefore requires dispersal of cells into fresh patches with replenished resources. We assume that this dispersal process occurs at fixed frequency of period *T*. The dynamics of the model then proceed in discrete generations (such as in a Wright-Fisher model (Blythe and McKane, 2007)), where a single generation consists of a growth phase followed by a dispersal phase.

**Figure 1:**
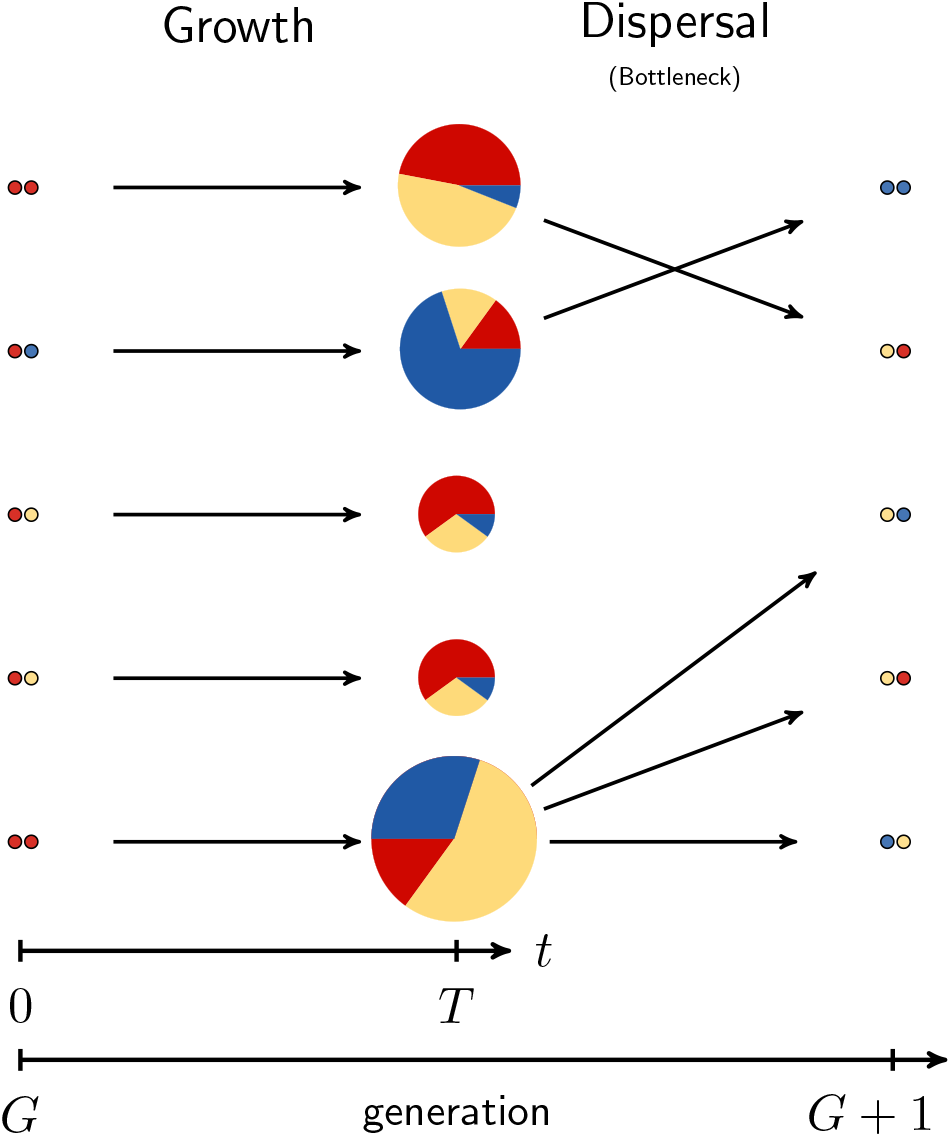
Overview of the model and dynamics over a single generation for a population of *M* = 5 patches. Each patch is colonised by a small number of cells at *t* = 0. A birth-death-mutation process then takes place (within-patch dynamics) over an interval of time [0, *T*]; different colours represent the growth rates of different cell types that have distinct growth rates. The pie charts represent the total population of each of *M* patches at dispersal. The size of slices in the pie charts represent the percentage of each type in the population and the overall size of the pies indicates the total relative population of the patch. After the growth phase, a dispersal event populates a fresh set of patches and in doing so creates a bottleneck, which in this example is a fixed size of 2.

Dispersal events also create bottlenecks in the process, hence each new patch is only colonised by a small number of cells. Dispersal is implemented as a random process such that larger patches (patches with a larger population of cells) are more likely to seed new patches. Thus selection at the level of patches favours patches comprised of many cells. Details of the two parts of the model, within-patch growth and dispersal, are given in the following sections.

### 2.1 Within-Patch Model

This part of the model describes the birth, death, and mutation of cells within a patch. Each patch initially contains resources that cells consume in order to reproduce. In reproducing, cells also create a waste by-product, with its accumulation contributing to the death rate of cells within each patch. To model these dynamics, we take an individual-based approach where we specify the states of the cells, the possible events and their rates (Black and McKane, 2012). The overall populations of cells, waste and resource within the patch at a given time are then specified by a continuous-time Markov chain (CTMC) (Black and McKane, 2012; Wilkinson, 2018). This implicitly assumes that the resource and waste are consumed and produced in discrete units. All quantities described below are relative to a single patch.

Cells are labelled according to their type, *i* = 1, …, *n*, where *n* is the maximum number of types tracked by the model. Cell types are distinguishable only by their growth rates, *β*_*i*_, hence we denote by *A*_*i*_(*t*) the number of cells of type *i* at time *t*. Similarly, we define *D*(*t*) and *E*(*t*) as the amount of waste and resource within each patch, respectively. The state of the system is then specified by the vector

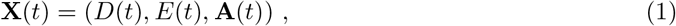

where **A**(*t*) is a vector with elements *A*_*i*_(*t*). To reproduce, cells pass through a cycle during which they consume *r* units of resource and produce *r* − 1 units of waste. The choice of *r* only affects the rate at which the resources are consumed, but the dynamics can be adjusted by scaling the initial resource, *E*(0) = *V* ≫ 1, to give similar growth trajectories regardless of *r*. Herein we set *r* = 4 and *V* = 10^6^.

Growth rates are discretised with mutation step size *μ*, where the growth rate of the *i*th type is defined as

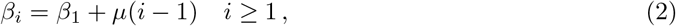

where *β*_1_ is the growth rate of the slowest growing type that is tracked by the model. Thus, the growth rate of type *i* + 1 is greater than the growth rate of type *i* and this ordering carries through to all elements of **A**(*t*). At each reproduction event, with probability *p*, instead of replicating to produce another cell of the same type, a mutation occurs to produce a cell of a different type. Mutations are modelled by changing the growth rate of the daughter cell by a single step either up or down. So if a mutation occurs to the daughter of a type *i* cell, with probability *q* the mutant will have a lower growth rate (*A*_*i−*1_ → *A*_*i−*1_ + 1), otherwise with probability 1 − *q* it is higher (*A*_*i*+1_ → *A*_*i*+1_ + 1). Thus *p* controls the overall probability of mutations relative to replication and *q* controls the symmetry of the process. This construction is similar to models of quasi-species (Eigen and Schuster, 1971; Nowak, 2006), but with a one-dimensional fitness landscape.

Within each patch we assume homogeneous mixing and hence the rates of events follows a mass action law that scales with the volume, *V* (van Kampen, 1992; Gillespie, 1977). The total rate of reproduction (including creating mutants) for type *i* is *V* ^*−*1^*β*_*i*_*A*_*i*_*E*, the product of the abundance of the cells, *A*_*i*_, the resource, *E*, and the growth rate, *β*_*i*_, divided by the volume. The rate at which cells of type *i* die is *V* ^*−*1^*A*_*i*_*D*. Hence the mean lifetime of a cell decreases as the amount of waste builds up in the patch and all cells die at the same rate independent of their type / growth rate. The events, transitions and rates for the model are summarised in Table 1.

**Table 1:**
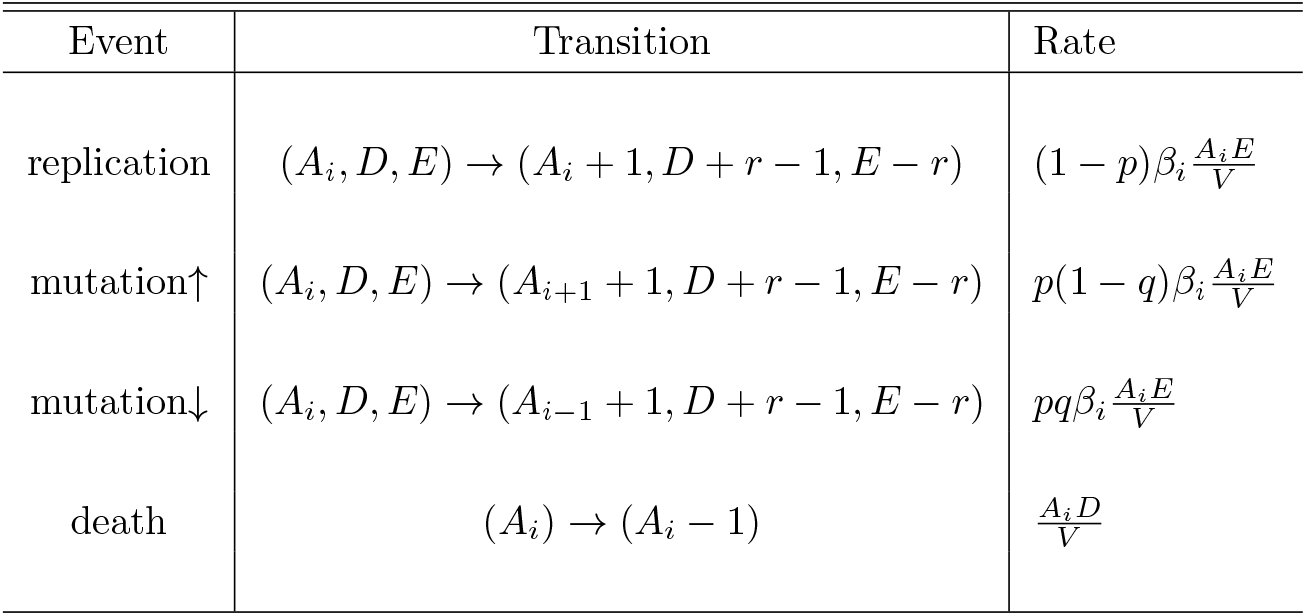
Events, transitions, rates, and propensity functions that define the within-patch model. Only components of the state that change in a given transition are shown, all other components are fixed.

| Event | Transition | Rate |
| --- | --- | --- |
| replication | $(A_i, D, E) \rightarrow (A_i + 1, D + r - 1, E - r)$ | $(1 - p)\beta_i \frac{A_i E}{V}$ |
| mutation↑ | $(A_i, D, E) \rightarrow (A_{i+1} + 1, D + r - 1, E - r)$ | $p(1 - q)\beta_i \frac{A_i E}{V}$ |
| mutation↓ | $(A_i, D, E) \rightarrow (A_{i-1} + 1, D + r - 1, E - r)$ | $pq\beta_i \frac{A_i E}{V}$ |
| death | $(A_i) \rightarrow (A_i - 1)$ | $\frac{A_i D}{V}$ |

Initial conditions for this process are

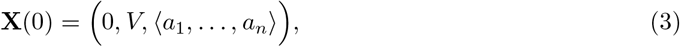

where *a*_*i*_ is the initial number of cell type *i* that colonise the patch, which is determined at the previous dispersal step. In specifying the initial conditions of patches, it is often simpler to label cell types by their growth rates rather than integers, where the mapping between type *i* and its growth rate *β*_*i*_ is given by Eq. (2). The initial condition for a patch with a bottleneck of size *b* can then be defined as a multiset (Knuth, 1997),

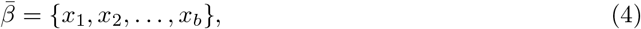

where *x*_*i*_ ∈ {*β*_1_, …, *β*_*n*_}. For example, {2.0, 2.0, 1.9} means a patch is initially colonised by two cells with growth rate 2.0 and one with rate 1.9. This can also be written more compactly as {2.0^2^, 1.9}.

### 2.2 Reordered State Vector Model

The model described above is a continuous-time Markov chain, hence trajectories can be simulated using the Gillespie (1977) algorithm, which generates exact sample paths of the process, or tau-leaping (Gillespie, 2001) which is a faster but approximate approach. Although in theory these algorithms can accommodate an infinite number of types, to make them computationally efficient we truncate the state space, i.e., set *β*_1_, the slowest growth rate tracked, and *n*, the total number of types tracked. These must be carefully chosen to avoid truncating the state-space to too small a region and it is not possible to do this a priori as the mean growth rate will evolve over generations of the model. Setting *n* too large affects the computational efficiency, which is important as the within-patch model must be run *M* times per generation for a large number of generations.

As will be demonstrated later, simulations of the model reveals that within a single patch it is only necessary to track a small range of growth rates, and the growth rates of the initial cells tend to be tightly clustered around a single value, even for much larger bottleneck sizes. This motivates the construction of a reordered state vector that takes advantage of the particular dynamics of our system and allows for a more more efficient simulation algorithm. We now define the growth rate of the *ℓ*th type as

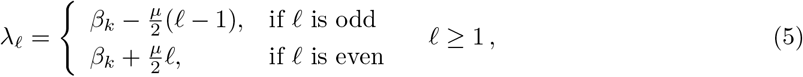

where *β*_*k*_ is chosen dynamically when the initial condition is set. This is chosen to correspond to the most populous cell type of the initial cells, which due to the dynamics of the model is usually close to the mean growth rate taken over the initial cells. If some cell numbers are equally populous then it is set to the fastest growth rate of this set.

This reordering essentially “folds” the state vector about *β*_*k*_ so that the bulk of the non-zero elements of **A**(*t*) remain in the first few entries of the vector. Similar ideas for re-ordering the state space to increase simulation efficiency have been proposed for a more general class of models (Cao et al., 2004; McCollum et al., 2006). The state space is still truncated by choosing a maximum value of *l* and altering the transition probabilities of the fastest and slowest types. Even with the dynamically updating *β*_*k*_, truncation means that it is still possible for initial cell growth rates to fall outside the tracked range. If this happens these cells are simply removed from the patch. This is an approximation, but in practice it works well as these cells are typically neither numerous, nor close to the mean growth rate of the remaining cells, and hence have a small impact on overall within-patch dynamics. Herein *l* is set at 23, which was found to be large enough, even for larger bottlenecks, such that truncation was rarely enforced.

Although the exact stochastic simulation algorithm is straightforward to implement, reactions occur frequently, which still makes the SSA impractically slow for large patch sizes. For this reason, the model is simulated using the standard tau-leaping algorithm (Gillespie, 2001), with interval of length *τ* = 0.1. This length is suitable as it retains the qualitative nature of the exact results, i.e., the general shape of the distribution of the total patch population at dispersal remains the same with this length of *τ*, but results in an algorithm that is more than 100 times faster than the SSA. More mathematical details of the re-ordering of the state vector are given in the Supplementary Material.

### 2.3 Dispersal

The second part of the model is a dispersal process that populates a new generation of patches and hence determines the initial conditions for the within-patch model. To specify this, we first define the matrix *A*, where each element *A*_*ij*_ is the number of cells of type *i* in patch *j* at the time of dispersal, *T*. Thus the total number of cells (of all types) in patch *j* is given by 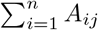. The dispersal process is simulated in two stages: first, *M* patches are sampled with replacement with probability in proportion to the total number of cells in the patch (i.e., the size of the patch),

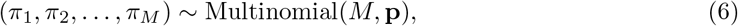

where the probability of sampling patch *j* is

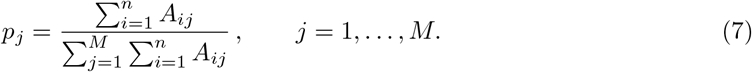

This gives rise to between-patch selection in the system as larger patches at the time of dispersal are more likely to seed future generations of patches. Next, we sample a bottleneck size, *b*_*j*_, and hence the number of cells to be sampled from each patch *π*_*j*_ as

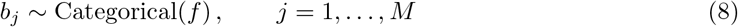

where *f* is the distribution over possible sizes. In many cases we take this distribution to be a delta function, *f*_*i*_ = *δ*_*i,B*_, so the bottleneck is of a deterministic size *B*.

Finally, to determine the initial conditions for the patches in the next generation we sample *b*_*j*_ cells from patch *π*_*j*_ in proportion to the overall number within the patch

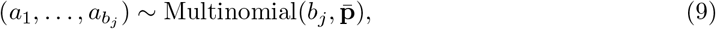

where 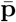 has elements

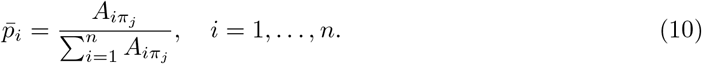

This procedure means that where the bottleneck is greater than one, all cells dispersed into a new patch come from the same parent patch.

### 2.4 Measuring evolutionary dynamics

We define a number of quantities that are useful in measuring the evolutionary dynamics of the system. The average cell growth rate within a patch at a given time *t* from the start of growth within that patch is calculated as,

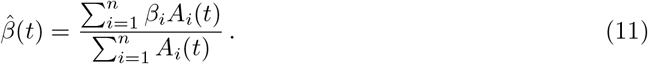

By tracking 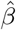 over growth and dispersal phases, it is possible to quantify the strength of selection at both levels of the model induced by competition and dispersal processes, respectively.

It is also informative to look at how different forces of selection evolve over generations of the model. This is done by examining how the average cell growth rate (defined in Eq. (11)) changes over the two phases that comprise a single generation of the model: patch growth and dispersal. The relative change in the average growth rate within a single patch in generation *k* is calculated as

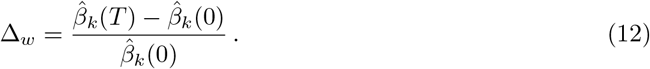

As defined, it is expect that this will be positive for our model because faster growing cells out compete slower growing cells within a patch, but only to a minor extent. The second quantity,

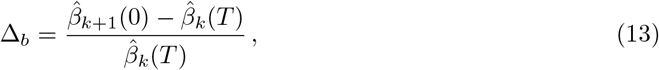

is the relative change in the mean cell growth rate between a given parent and offspring patch. The term 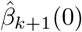 is the average cell growth rate within a patch after dispersal has taken place. On average, we expect this to be smaller than the growth rate within a patch immediately before dispersal, 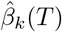, since dispersal favours slower growing cells. As defined, this means Δ_*b*_ should be negative. Note that with these quantities defined, the average growth rate per generation can be decomposed as the sum of the within- (Δ_*w*_) and between-patch (Δ_*b*_) forces

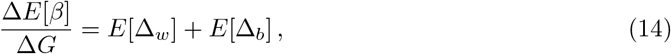

where the expectations are over the population of patches in the system.

## 3 Single Cell Bottleneck Dynamics

In this section we discuss the dynamics of the model when it is assumed that the dispersal process imposes a strict bottleneck and hence each patch is founded by a single cell (*b* = 1). This allows connection to the previous work of Black et al. (2020) and establishes results that can be usefully compared to those generated in the following section where this assumption is relaxed.

Figure 2(a) shows the total population within a patch for five independent realisations of the model, each founded by a single cell with the same growth rate 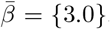. Initially when the number of cells is small there is a phase where the dynamics “stutter” but this lessens once populations grow large enough and exponential growth begins (Black et al., 2014). During growth, resources are depleted, which slows the growth rate, and waste accumulates increasing the death rate of cells. At some point the number of births and deaths balance and populations peak and then decline. The rate of cell death is proportional to the amount of waste accumulated in the patch. As this waste is initially zero, there is essentially no chance that the population of cells dies in the initial stochastic growth phase.

**Figure 2:**
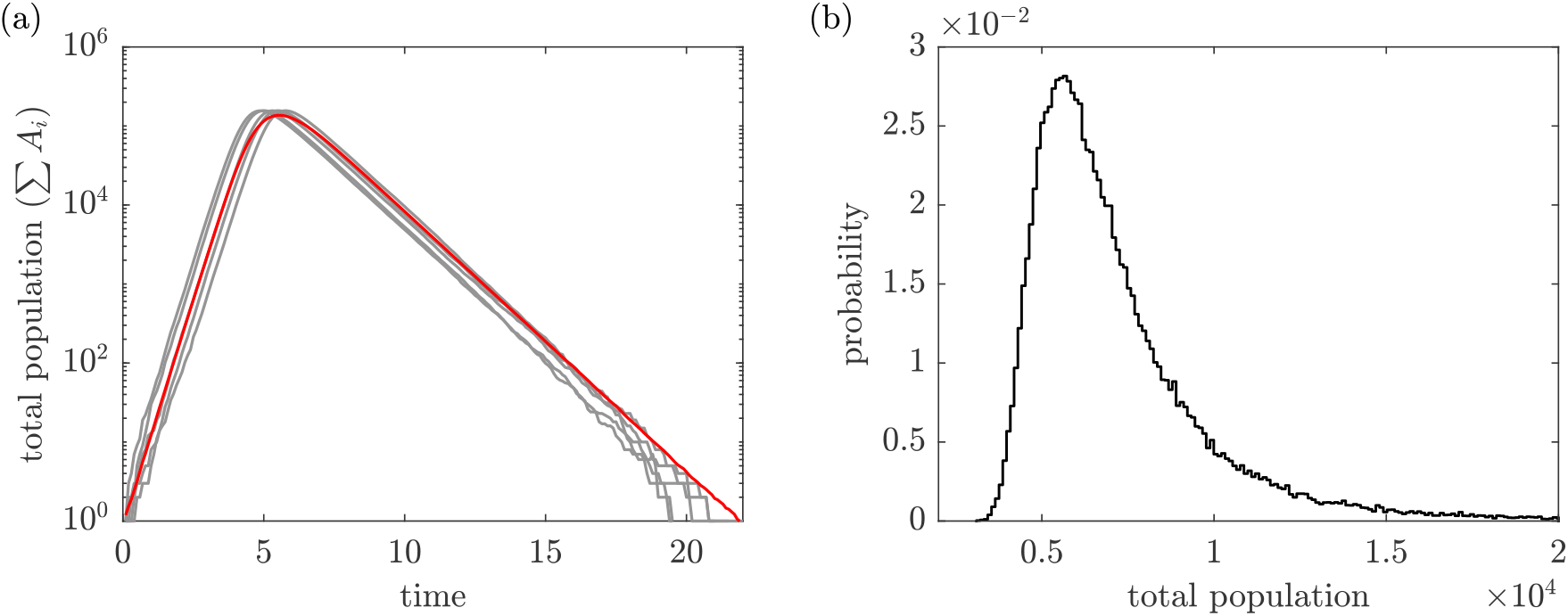
Trajectories of cell populations within a patch. (a) Five realisations of the total patch population as a function of time (grey lines) and the mean total population (red line). Patch population is computed as the sum of individual cells in a patch. (b) The total size distribution estimated at *t* = 10 from 10^5^ realisations. Other parameters: *V* = 10^6^, 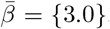, *p* = 0.01, *q* = 0.5, *μ* = 0.1.

The initial stochasticity affects the time until exponential growth is reached and hence the time for populations to peak. This translates into variation in the size of the patches at any given fixed time after seeding. This is illustrated in Figure 2(b), which shows the patch size distribution at *t* = 10. In this example, where patches are sampled a time long after the populations have peaked, the distribution is skewed with a heavy tail that leads to a higher mean, relative to the mode. This is a consequence of some cells entering exponential growth later than others (due to stochastic fluctuations) and so also entering the decline phase later. This natural variability in the patch size was not present to such an extent in the previous model of this process, but was possible to introduce with the addition of extra variability in the dispersal time (Black et al., 2020).

Figure 3(a) further illustrates the patch sizes and composition for 64 realisations of the within-patch model at *T* = 10 founded by single cells with the same growth rate, 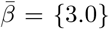. Figure 3(b) shows, for a single realisation, the populations of the cells by their type, *A*_*i*_(*t*), as a function of time. Together these results show that mutants of the original type are not produced until after the original type has reached an appreciable level and mutants of mutants (second order mutants) are comparatively less common. Small bottleneck size at dispersal thus imposes strong homogeneity on the composition of patches at later times. A further point concerning the dynamics of the patches that is important: after the population has peaked, the proportion of different types within each patch remains largely fixed. This is a consequence of the rate of cell death, which is identical for all cells (see Table 1 for the model rates).

**Figure 3:**
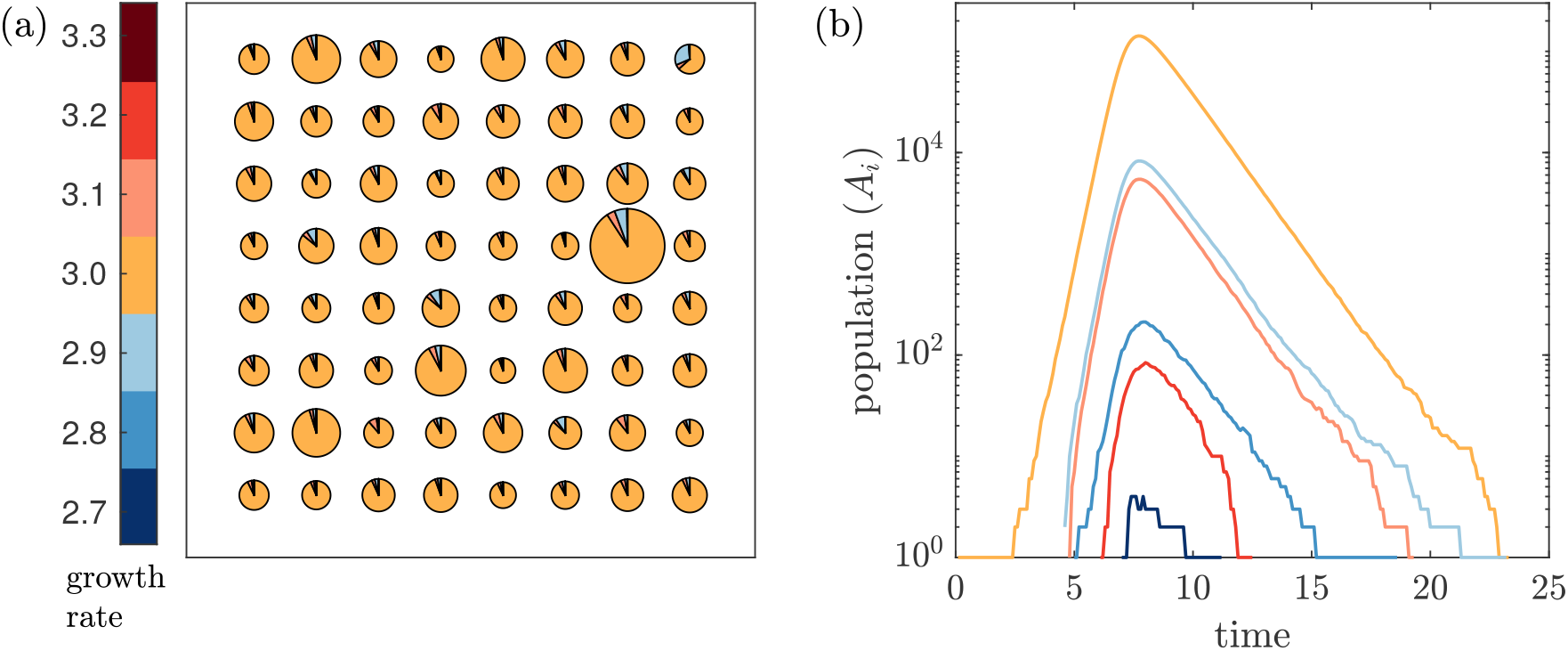
Illustration of patch sizes and composition at dispersal. (a) 64 realisations of the within-patch model with the same initial condition, 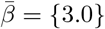, at dispersal time *T* = 10. Each pie has area proportional to the total number of cells with arcs proportional to the composition. (b) A single realisation showing populations by type as a function of time over the growth phase. Both the pie charts and lines are coloured according to the growth rates for each cell type. Parameters are as in Figure 2.

Figure 4(a) and (b) shows the time resolved dynamics of a number of independent realisations of the full evolutionary model for two different dispersal times, *T* = 4 and 10. The dispersal times are chosen to be short and long, respectively, compared with the time for the population to peak with an initial growth rate of 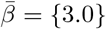 (see Figure 2(a)). In both scenarios, the fitness of patches (reflected in the population size of cells within patches) increases before reaching an equilibrium and this is achieved through changes in cell fitness (cell growth rate). When *T* is short, the forces of selection within the patch and in the population of patches are aligned (faster growth rates results in both fitter cells and patches). When the dispersal time is long, the growth rate of the cells decreases over generations of the model. This phenomenon has previously been called “fitness decoupling”, referring to the fact that fitness at the two levels is no longer aligned (Michod, 1999; Okasha, 2008; Black et al., 2020; Bourrat, 2021).

**Figure 4:**
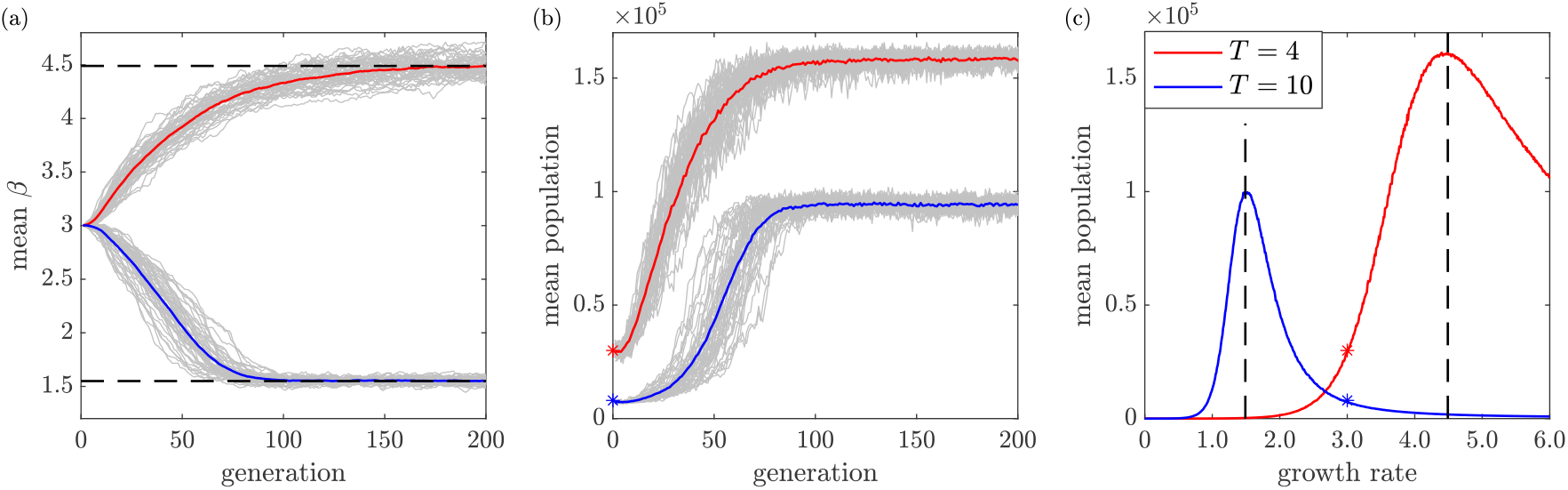
Evolutionary dynamics for a system with a bottleneck of one cell, and for two different dispersal times, *T* = 4 and 10. (a) The average growth rate over a population of 100 patches and (b) the average total population per patch. The grey lines show 50 individual realisations, and the coloured lines show the averages over these realisations. In both cases, the dispersal process creates selection pressure that favours larger patches at the time of dispersal. Dashed lines show the equilibrium growth rates and the stared points represent patch population sizes for the initial growth rate of the simulations. (c) Fitness landscape view of the evolutionary process for fixed dispersal times *T* = 4 and 10. Each curve shows the average total population within a patch for a fixed dispersal time as a function of the growth rate of the initial cell that colonises a patch. When *T* = 4, the population peaks at a growth rate of *β* ≈ 4.5 and when *T* = 10 the maximum population is reached when *β* ≈ 1.5. These peaks, indicated by the dashed lines, correspond to the equilibrium growth rates reached by the simulations shown in (a). The stared points indicate the patch population sizes for the initial growth rate of the simulations as seen in (b).

With a single cell bottleneck, the evolutionary outcomes and dynamics of the system can be understood using a simple fitness landscape approach (Nowak et al., 2010). This is valid because the accumulation of mutants in a patch is small and hence the *mean total population* of the patch at the time of dispersal is strongly correlated with the growth rate of the founding cells. Landscapes are derived for given values of *T* by computing the mean total population as a function of the growth rate of the initial founding cell; the resulting curves are plotted in Figure 4(c). These show peaks at different positions for different dispersal times, which represent equilibrium points of the evolutionary dynamics. If the model is initialised such that the growth rate of the cells is away from the equilibrium, then these curves indicate that we will observe evolution ‘up the hill’ towards the peaks (as seen in Figure 4(a)). The peaks represent the cell growth rate that optimises the size of the patches on average for a given dispersal time. Figure 4(c) shows that it is the relative length of the dispersal time compared to the time to peak for an initial growth rate that is important in setting the direction of evolution in these populations. For example, fixing *T* = 10, if the cell growth rate was initialised at values *<* 1.5, then we would instead observe an increase in the growth rate. More results for the evolutionary dynamics with a single-cell bottleneck, and how these change with mutation rate, *μ*, mutation probabilities, *p* and *q*, and the number of patches in the system, *M*, are discussed in the Supplementary Material.

Ecological conditions are key to understanding the evolutionary dynamics of these nested populations when the dispersal time is long. Limited resources within a patch restrict the total number of cell divisions possible. This, coupled with the accumulation of waste, which rapidly leads to cell death after a certain point in time, effectively limits the period of time over which cells can reproduce and hence also limits the production of mutants. This can be contrasted with, for example, growth in a chemostat where resources are constantly supplied, and waste removed. In such a reactor, faster growing cells have time to out-compete slower types and drive them to extinction. This is not possible in our model ecology, as even though slower growing mutants are at a disadvantage within the patch, there is insufficient time for them to be driven extinct (excluding when the dispersal time is so long that all cells go extinct). Thus, patch ecology allows for the production of a small but significant fraction of slower growing types that can be dispersed.

In our model, the bottleneck has two functions. Within a patch, it suppresses the growth of mutants (both faster- and slower-growing) and hence competition. At the population level, it promotes patch fitness at the expense of the average cell growth rate within the patch: if a slower growing type is by chance dispersed, then it is itself initially free from competition in a patch, which allows it to become established. In contrast, when the dispersal time is short, cell and patch finesses are aligned (faster growing cells lead to larger patches) and hence this property of the bottleneck is not important. For the remainder of this paper, we concern ourselves only with the situation in which the dispersal time is long compared with the time to reach peak population for an initial cell growth rate, and hence increased patch fitness is achieved by decreasing cell growth rate.

## 4 Multi-cell bottleneck dynamics

In the previous section, we showed how a bottleneck of one cell facilitates an evolutionary process where between-patch selection created by the dispersal process dominates within-patch selection (for higher growth rates) leading to an overall reduction in the average growth rate of cells over a number of patch generations. We now expand these results to investigate how the evolutionary process changes when larger bottlenecks are allowed. Simulations are initialised from a homogeneous population of cells with the same growth rate, i.e., for each patch **X**(0) = (0, *V*, ⟨0, …, *a*_*i*_, …, 0⟩), where *a*_*i*_ is the number of type *i* cells. In terms of the multiset notation defined in Section 2.1 this initial condition is written as 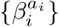. We initially describe results for smaller values of *b* before moving to larger values where the picture becomes more complex.

Figure 5 shows evolution in mean growth rate, patch size and within- and between-patch selection averaged over realisations of the process for a series of increasing bottleneck sizes, *b* = 1, …, 5. For these sizes, as the system moves through generations, the composition of patches at the time of dispersal shifts toward cells with slower growth rates and the average patch size increases before eventually reaching equilibrium. We can observe two main effects of increasing the bottleneck size: a slowing-down of the evolutionary dynamics and also a change in the equilibrium values of both the mean growth rate and patch size.

**Figure 5:**
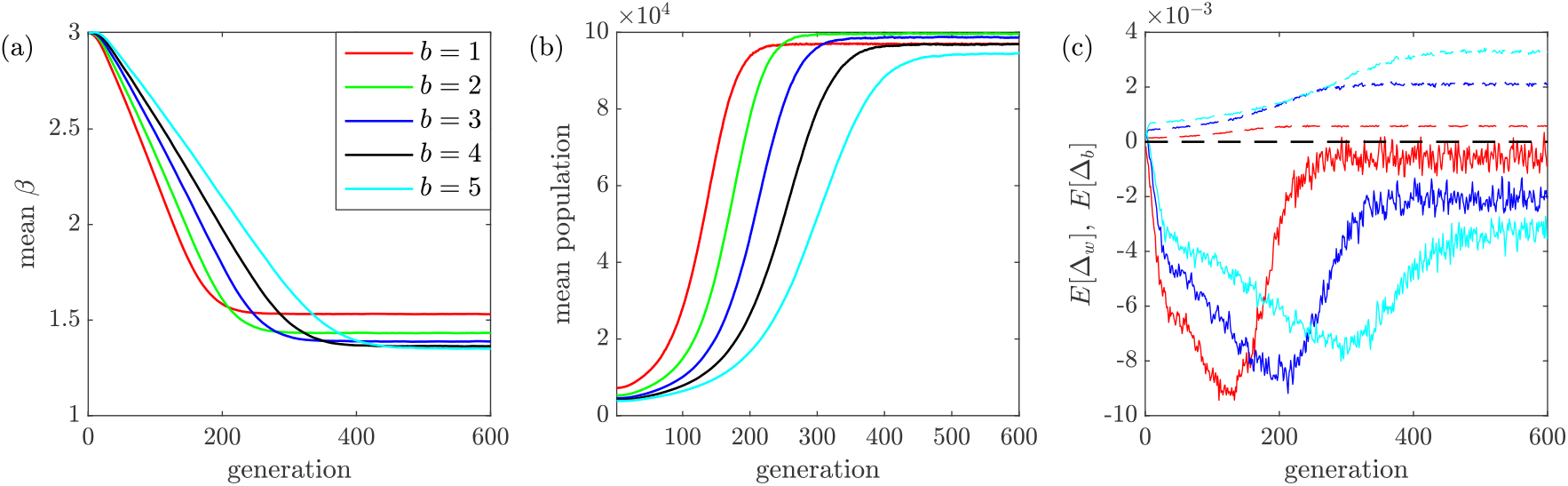
Evolutionary dynamics for a system with varying bottleneck sizes. (a) The average growth rate, (b) the average total population per patch and (c) the average change in growth rate decomposed into within- (dashed lines) and between-patch (solid lines) components. Each line shows the averages over 50 individual realizations. Parameters: *μ* = 0.05, *p* = 0.01, *q* = 0.5, *T* = 10, *M* = 100.

The decrease in rate of change in average patch size and growth rate can be attributed to changes in the forces of selection at both levels of the model. These forces can be quantified by the expressions given in equations (12) and (13), which measure how the growth rate changes over a single growth phase (Δ_*w*_), and after a dispersal phase (Δ_*b*_), respectively. The changes in the expected values of these two quantities, averaged over patches and realisations are shown in Figure 5(c). The sum of these two quantities corresponds to the rate of change (the first derivative) of the curves shown in Figure 5(a).

Larger bottlenecks increase within-patch selection as demonstrated by dashed curves shown in Figure 5(c) that are shifted upward for lager values of *b*. This is an obvious implication of the additional competition between cell types within a patch. With larger bottleneck sizes, competition is present from the beginning of patch colonisation, rather than arising from mutations later when *b* = 1. However, because the growth phase is time-limited, cell types do not in general go extinct and this still restricts competition between the types to some extent. Larger bottlenecks also decrease between-patch selection, as can be observed by the solid curves in Figure 5(c). This is because larger bottlenecks lower the fidelity of transmission of phenotype at the patch level (where the phenotype is patch size). With a larger bottleneck, different configurations of initial cell growth rates tend to have similar size distributions at the time of dispersal. Since patches are chosen in proportion to their size at dispersal, similar size distributions result in a weaker force of selection.

This effect can be observed by contrasting some patch size distributions for a single cell and three-cell bottleneck as shown in Figure 6. For the three-cell bottleneck, both in and out of equilibrium, the correlation between the growth rate of the initial cells and the mean patch size is weakened, and hence on average the difference in final size between similar configurations is smaller. For even larger bottlenecks, this effect is magnified due to the combinatorial explosion in the possible initial configuration of cell growth rates. Another way of conceptualising the weakening of the phenotype connection is through the effect on the fitness landscape. The multiplicity in the possible cell combinations means that the fitness landscape grows exponentially in dimension (and hence is difficult to visualise) and becomes flattened (Reidys and Stadler, 2002). The increase in dimension means evolution has to proceed by smaller steps in an absolute sense, and the flattening means the fitness difference between these steps is smaller.

**Figure 6:**
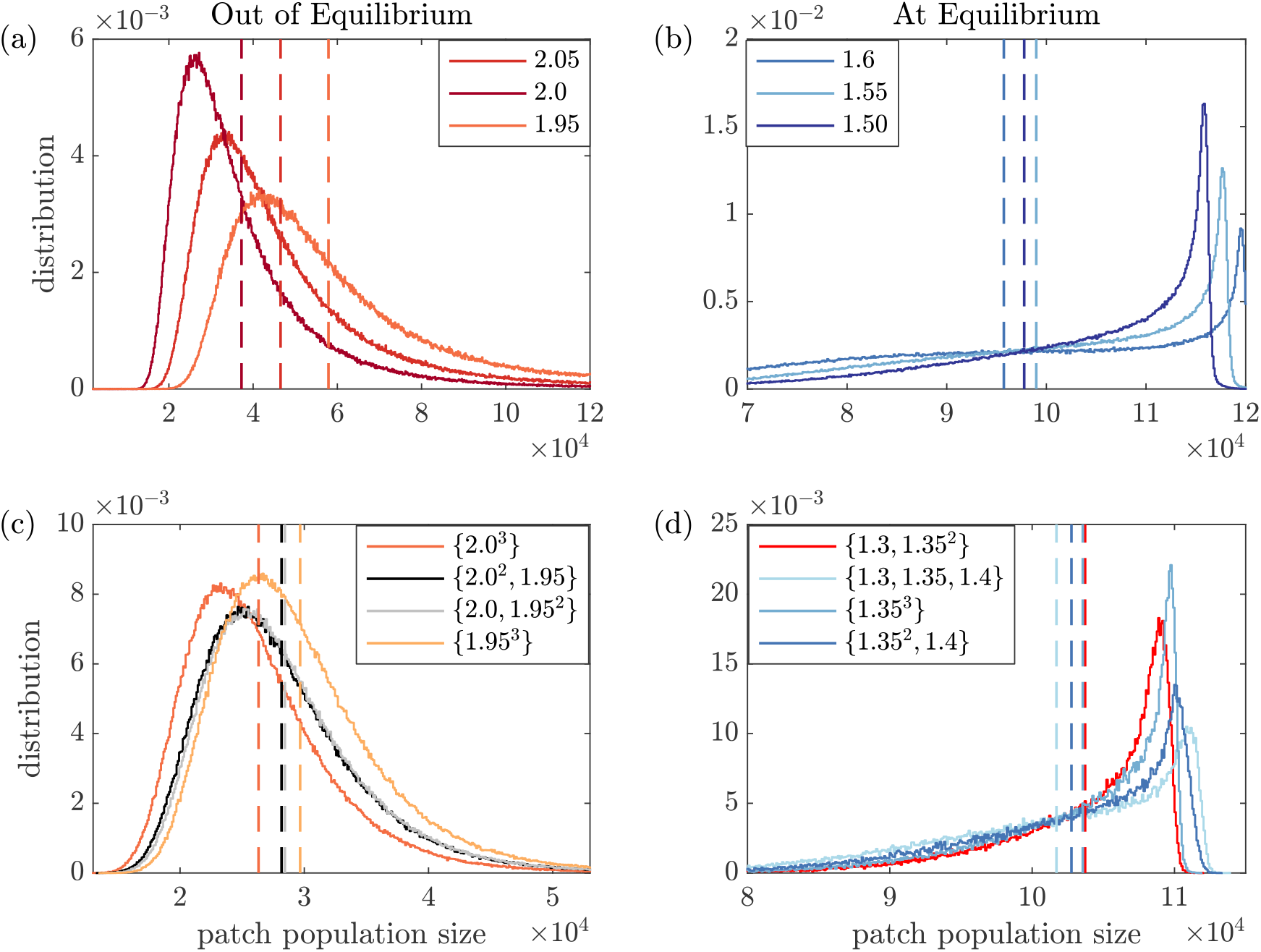
Patch size distributions at the time of dispersal (*T* = 10) for different initial growth rates and bottleneck sizes, both in an out of equilibrium. Panel (a) shows results for a single cell bottleneck out of equilibrium, and (b) at equilibrium (*β* ≈ 1.55 in this case). Panels (c) and (d) show the same quantities for a three-cell bottleneck. Note for *b* = 3 the equilibrium growth rate is lower (*β* ≈ 1.35). In all panels, vertical dashed lines indicate the mean patch size for each distribution.

The small changes in the equilibrium growth rate visible in Figure 5(a) and (b) are due to a combination of changes in the forces of selection as well as changes in the within-patch dynamics resulting from the population starting at a larger initial size. These distort each other resulting in non-monotonic behavior of the equilibrium as the bottleneck is increased. This is discussed in more detail in Section 4.1.

Figure 7 shows simulations for larger fixed bottleneck sizes of 10, 15 and 20. As *b* increases, a further slow-down of the evolutionary dynamics is observed, along with a rise in the equilibrium growth rate, before an eventual reversal, where the growth rate no longer decreases, but instead increases without bound. This is conceptually similar to a phase transition through a critical point (Stanley, 1987). Around *b* = 15, the forces of selection within the patches and between the patches are almost equal and critical slowing down is observed (Elf et al., 2003), where the size of fluctuations becomes very large. After this point within-patch selection dominates between-patch selection leading to the continual rise in cell growth rate over generations.

**Figure 7:**
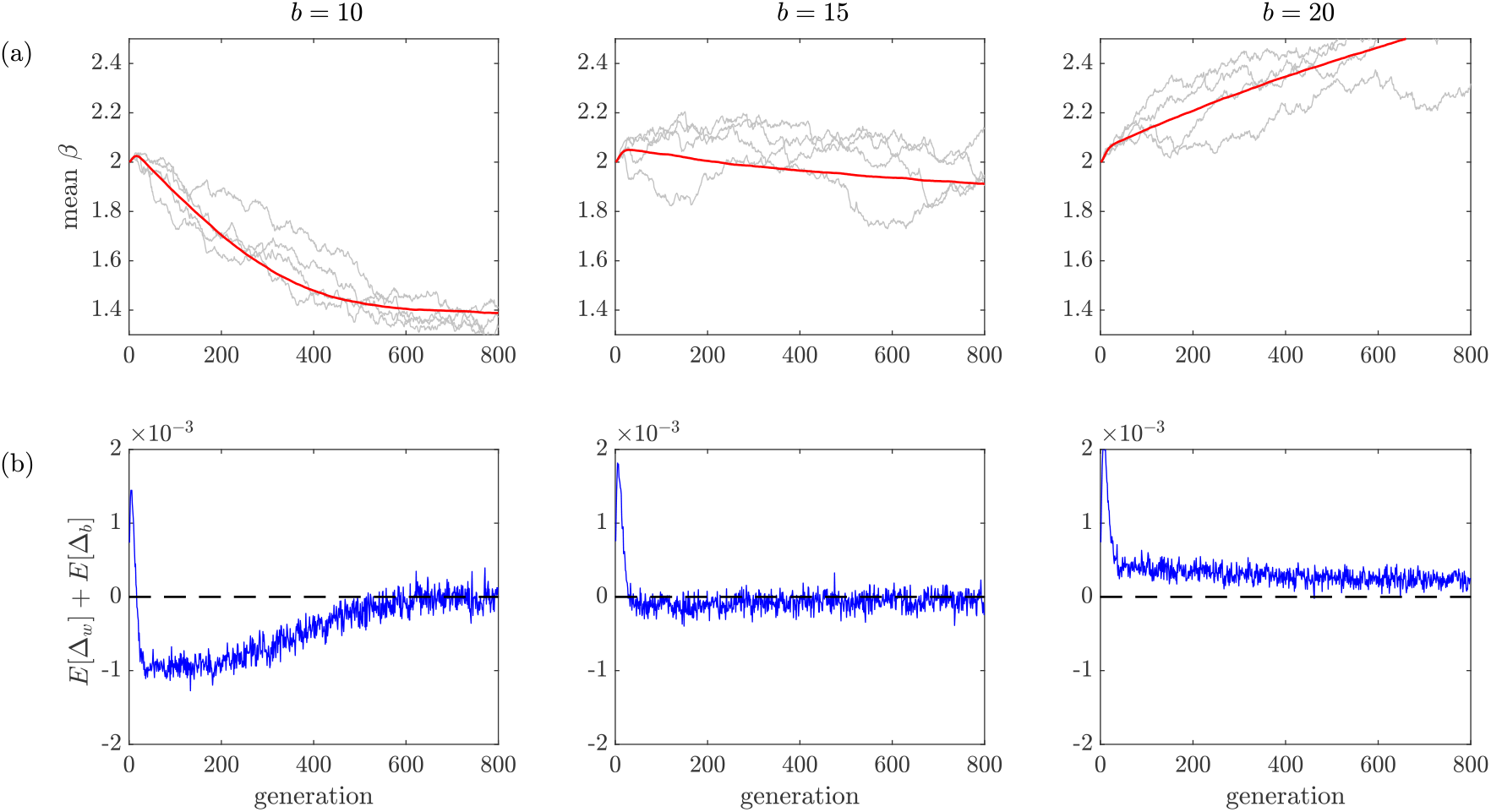
Evolutionary dynamics with larger bottlenecks. (a) Change in the average growth rate over generations. (b) The sum of the within- (Δ_*w*_) and between-patch (Δ_*b*_) forces of selection, which illustrates the net force of selection (net change in the growth rate per generation, see Eq. (14)). For *b* = 10 and 15, simulations eventually reach equilibrium hence the difference goes to zero. For *b* = 20, within-patch selection always dominates between-patch selection and so the difference remains positive. Model parameters: *M* = 100 patches, *μ* = 0.1, *p* = 0.01 and *q* = 0.5.

### 4.1 Equilibrium behavior at larger bottlenecks

A complex pattern emerges in equilibrium values with increasingly large bottlenecks. This is further complicated by the the possibility of reversal in the magnitudes of the two forces of selection, as described above, and hence failure of the system to reach equilibrium. Figure 8 shows the mean and variance of the growth rate and patch size, both at equilibrium, as a function of bottleneck size. We run the same analysis for *M* = 100 and 500 patches. A larger number of patches reduces the stochasticity resulting from the dispersal process (see Supplementary Material) hence between-patch selection is stronger for *M* = 500. Note that the blue curve does not continue past *b* = 15 as an equilibrium is no longer reached in this case.

**Figure 8:**
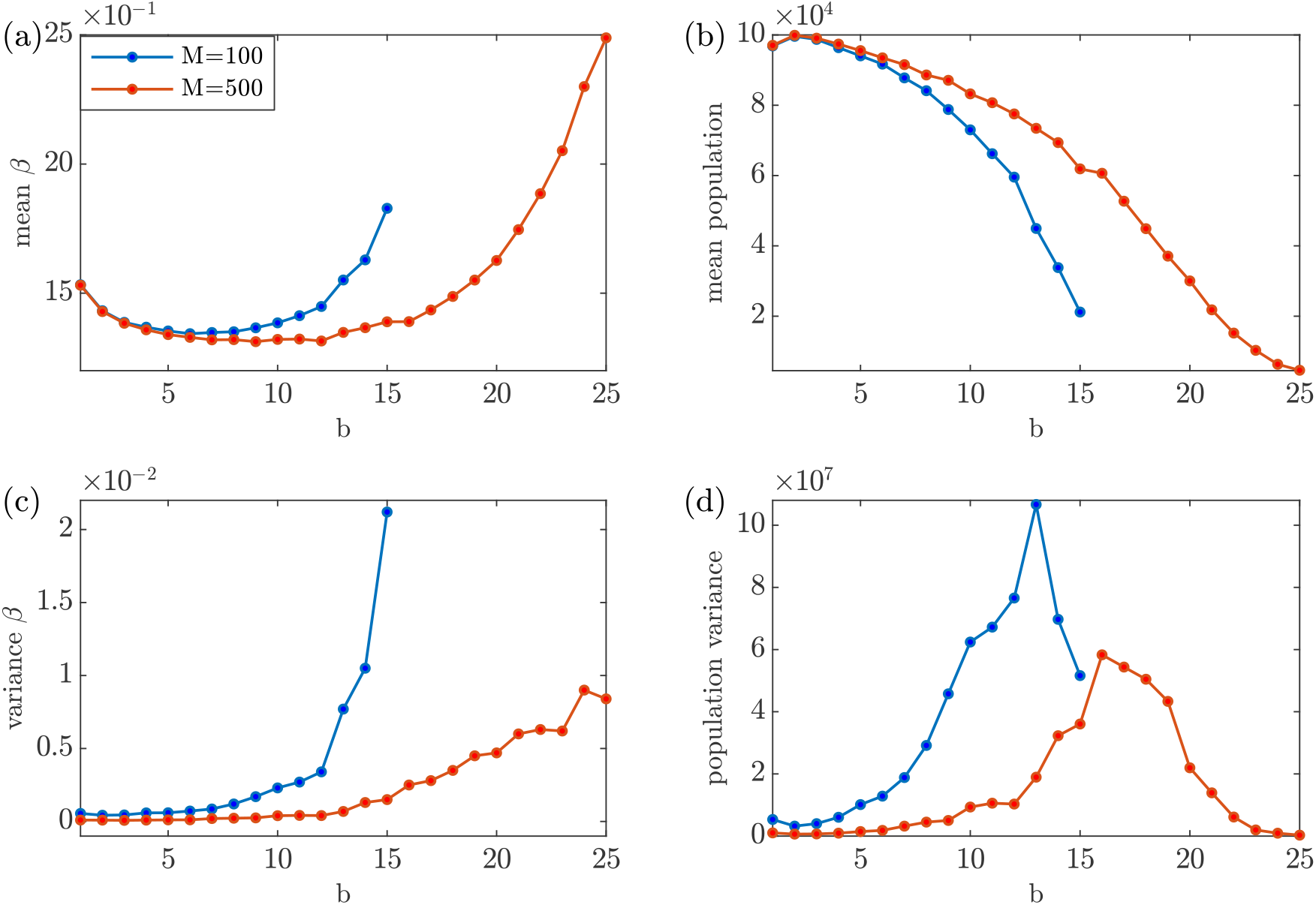
The mean and variance of growth rate and patch size at equilibrium as a function of bottleneck size for *M* = 100 and *M* = 500. (a) and (c) show the mean and variance in the cell growth rate. (b) and (d) show the same for the patch size. Note that the curve for *M* = 100 does not continue after *b* = 15 as there is no longer an equilibrium (see Figure 7). Other parameters: *μ* = 0.05, *q* = 0.5, *p* = 0.01 and *T* = 10.

At *b >* 2, the mean patch size (Figure 8(b)) shows a steady decrease. This is due to within-patch dynamics that change as the initial number of cells grows. Larger bottlenecks means that patches are founded by more cells, but additionally, populations enter exponential growth phase more rapidly. This, in turn, means that a larger bottleneck reduces the time for the population to peak on average (see Figures S8 and S9 in the Supplementary Material). Hence for the same dispersal time, larger bottlenecks result in a smaller patch size on average.

The variance in patch size initially increases before decreasing. The initial increase is due to the changes in the forces of selection and the increasing number of configurations resulting in a larger range of patch sizes at equilibrium. The decrease at even larger sizes is because the patches can become so small that extinction becomes possible. In the patch dynamics, a population of zero is an absorbing state and this skews the size distribution.

The mean growth rate initially drops as the bottleneck increases in size before rising. The initial decrease is due to changing of the peak time with bottleneck size as discussed earlier. Larger bottlenecks result in a smaller patch size on average and so the equilibrium growth rate to optimise the patch size tends to decrease in order to compensate. This is eventually counteracted by increasing within-patch selection leading to the later rise.

The sharp rise in the variance in the growth rate for the *M* = 100 case is clearly seen (Figure 8(c)), indicating the position of the reversal in the magnitude of the forces of selection. It is interesting that, at least for these parameters, with enough patches in the system, between-patch selection is always large enough to curtail increased within-patch selection at larger *b*.

## 5 Random Bottleneck Dynamics

Up to this point, we have assumed that the bottleneck is a fixed, deterministic, size across all patches and generations. We now investigate changes in dynamics when this condition is relaxed and bottleneck sizes are randomly sampled. Each time a patch liberates dispersing cells, the number of migrating cells is drawn at random from a fixed distribution (see Eq. (8)), where *f* = (*f*_*i*_)_*i*=1:*N*_ is the distribution over the possible sizes, and *f*_*i*_ is the probability of size *i*.

Figure 9 compares evolution of the mean growth rate for several bottleneck size distributions. In each case, fixed bottleneck size dynamics are compared to bottleneck sizes chosen from a discrete distribution, where 𝒰 (*A*) is uniform over the finite set *A*. In Figures 9(a) and (b) that include only smaller bottlenecks, we see that the dynamics with a uniform distribution over the sizes lies between the fixed-size extremes. That is, when bottleneck size is drawn from a uniform distribution, the mean growth rate remains between the corresponding maximum and minimum fixed bottleneck sizes in that distribution. Even when the bottlenecks are larger (Figure 9(c)), where the growth rate would be expected to rise if it was a fixed size, we instead see that the mean growth rate drops, implying that within-patch selection is sufficiently curtailed by the presence of single-cell bottleneck events. This suggests that only occasional small bottleneck events are necessary for significant evolutionary change in the growth rate of this system.

**Figure 9:**
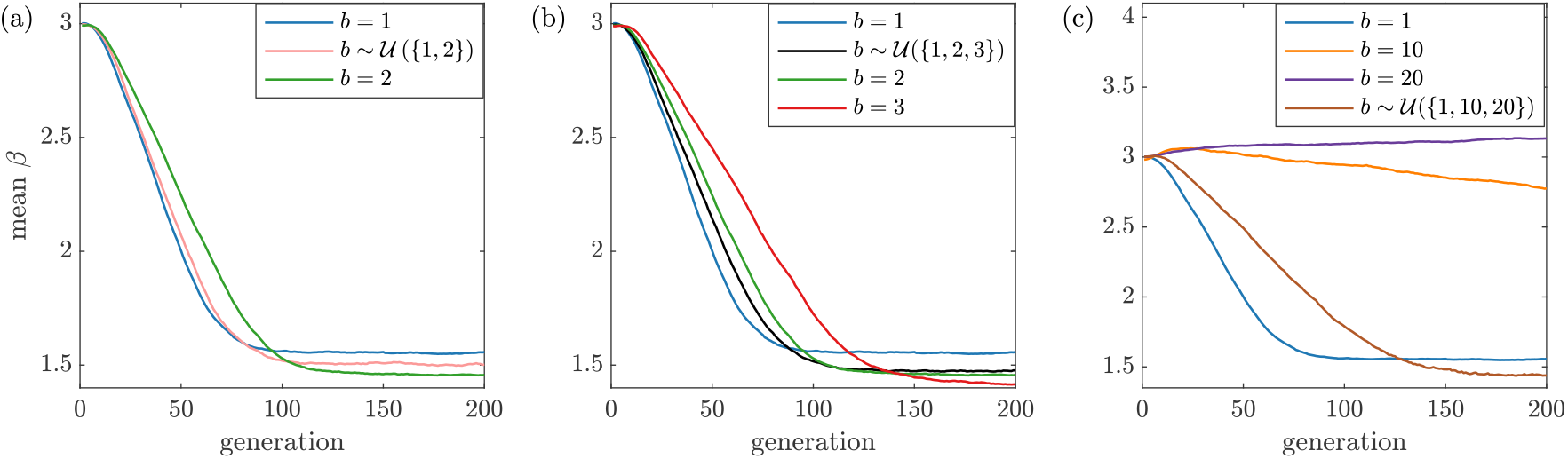
The evolution of average growth rate when bottleneck size is chosen from a discrete distribution, where 𝒰 (*A*) is uniform over the set *A*. (a) and (b) show the dynamics for mixtures of smaller bottlenecks, (c) shows the effects of a distribution where the size is either large or unicellular.

We further investigate these effects by focusing on a system with only two possible bottleneck sizes: one and twenty, and changing the probability distribution over these sizes. The results of this investigation, shown in Figure 10, demonstrate that only a few patches with a single-cell bottleneck are required per generation to suppress within-patch selection and generate faster evolutionary change in growth rate. Thus, even if a bottleneck of size one occurs for only five patches out of 100 in a generation, a decrease in the average growth rate is observed. This is initially surprising, but can be explained by the homogenising effect that smaller bottlenecks have on the composition of patches, as illustrated in Figure 11. The effect, once initiated, persists through subsequent generations, even though later bottlenecks are much larger. This homogenisation also means that the phenotypic link between the initial composition and the patch size is temporarily strengthened and so selection will have a greater effect at the higher level. That is, a patch colonised by 10 cells of the same type will grow to a more deterministic range of sizes than a patch composed of 10 different types.

**Figure 10:**
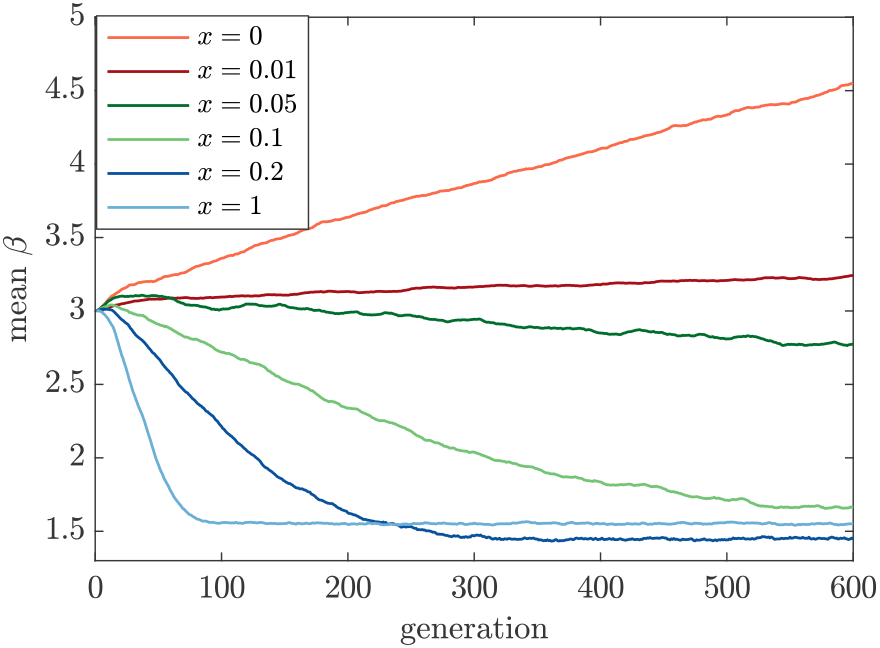
Evolution in the mean growth rate, *β*, for a non-uniform, binary bottleneck distribution. Bottleneck sizes are drawn from a distribution where *f*_1_ = *x* and *f*_20_ = 1 − *x* with 0 ≤ *x* ≤ 1. Thus *x* = 1 corresponds to a strict unicellular bottleneck (*b* = 1) and *x* = 0 a bottleneck of *b* = 20.

**Figure 11:**
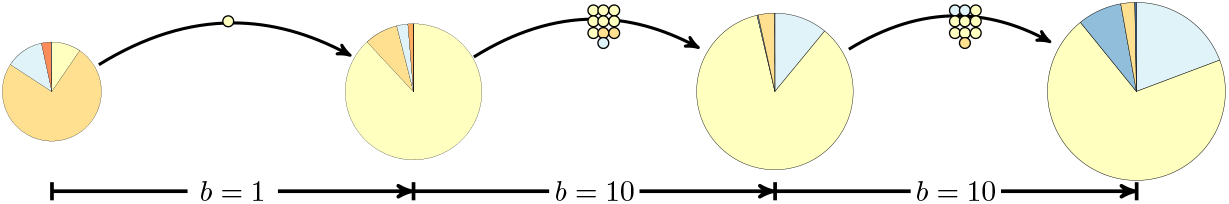
Illustration of the homogenising effect of a randomly small bottleneck. If by chance a single slower-growing cell is dispersed, then this gains a larger advantage by freeing it from competition and this propagates through many subsequent generations.

## 6 Discussion

Black et al. (2020) showed that simple environments consisting of finite patchily distributed resources and a periodic dispersal process that also imposes a bottleneck can scaffold a multi-level Darwinian process. When the period of time between dispersal events is long, between-patch selection causes patches to increase their fitness while driving a reduction in cell growth rate. This tradeoff creates suitable conditions for the first steps in the transition from unicellular to multi-cellular life. Building on this previous work, we have developed a fully stochastic model of an ecologically scaffolded population to further investigate the role of bottleneck size on the evolutionary dynamics of this system. The model presented in Black et al. (2020) was only partially stochastic, capturing the effects of mutations and stochastic dispersal times, but did not naturally allow for bottlenecks larger than one due to certain approximations used to derive the dynamics.

In common with other studies on the role of bottlenecks (Michod and Roze, 1999; Bergstrom et al., 1999; Chuang et al., 2009; Cremer et al., 2012; Melbinger et al., 2015; Rose et al., 2020; Doulcier et al., 2020), we see that the bottleneck mediates opposing selection forces, over different timescales, within and between patches. In contrast to others, we have adopted a mechanistic modelling approach, so the dynamics are a function of the interaction of the different components rather than simply being imposed phenomenologically. This has allowed a fine-grained investigation of how the bottleneck affects the forces of selection at both levels of the model by *measuring* its effect in silico. To the best of our knowledge, this is the first analysis of a dynamic bottleneck with an emphasis on stochastic effects and few cells, which has so far been absent from the literature. A drawback of our approach is that we can only simulate our model and hence cannot derive analytic results, but the complexity of the process and the multi-scale nature of the problem precludes this for now.

Concentrating on the region of parameter space where we see a decrease in cell growth rate with a single-cell bottleneck, we find that the process is relatively insensitive to larger bottlenecks up to a point before the size becomes too big and within-patch selection, caused by competition between cells with different growth rates, dominates between-patch selection arising from the dispersal process. Surprisingly, allowing for random bottleneck size distributions did not significantly change the evolutionary dynamics compared to the deterministic-size case and in fact enhanced the suppression of within patch selection more than would be naively thought. Thus, in a system with a normally large bottleneck that would result in limited evolutionary change in patch size and cell growth rate, these dynamics can be accelerated by adding only infrequent small bottlenecks. This finding enhances the robustness of ecological scaffolding as an explanation for the evolutionary transition to multicellularity as it demonstrates that a very strict bottleneck is not a necessary condition for an ETI to be initiated, infrequent smaller bottlenecks may be enough.

This result brings into question the abundance of single-cell bottlenecks in nature. Indeed, if a single-celled bottleneck is not required for ETIs to be initiated, then why are they so ubiquitous in nature? We have observed that single-cell bottlenecks create stronger between-patch selection and hence generate faster evolutionary dynamics. It might be that bottleneck size is itself an evolved trait. Such a possibility seems highly likely in the face of an innovation that generates a reproductive division of labour: a lineage that has evolved a division of labour between, for example, soma- and germ-like cells will be out-competed by a lineage that lacks such a division in cases where patches are established by cells of the two types. Additionally, more complex developmental processes, which we have not attempted to model here, would likely benefit from smaller bottlenecks.

Attempts were made to use the model presented herein to additionally investigate the evolution of bottleneck size. This builds on the random bottleneck model presented in Section 5. In the results presented, the parameter *x* controlling the probability of a larger or smaller bottleneck is fixed, but this can also be made an intrinsic property of the cells. So when dispersal occurs, a single cell is initially selected and its value of the parameter *x* is used to randomly determine if more cells are dispersed along with it. Thus, *x* can be thought of as the ‘stickiness’ of the cell and allowed to evolve along with growth rate. Extending the model in this way adds considerably to its complexity as two traits now need to be tracked for each cell. Preliminary results indicate that faster evolutionary change as a result of smaller bottleneck sizes can drive evolution in the ‘stickiness’ trait to some extent. However, pinning down the actual cause of this has proved difficult and it cannot be ruled out that the observed results are an artifact of other effects of changing the bottleneck size, such as on the final size of the patch as discussed in Section 4.1. A thorough investigation of the evolution of bottleneck size likely requires a different model where the final size of patches is less sensitive to the size of the bottleneck, which is not achievable with the current model.

Although this work was first conceived to understand evolutionary transitions in individuality, parallels and insights into the evolutionary dynamics of host-pathogen systems are obvious. Nested models have been successful in emphasising the importance of within-host disease dynamics on pathogen evolution for some time (Gilchrist and Coombs, 2006; Alizon and van Baalen, 2008; Luciani and Alizon, 2009; Saenz and Bonhoeffer, 2013). These consist of both essential and inessential nested hierarchies which are differentiated by the extent to which feedback between levels is incorporated. The standard approach is to model within-host dynamics with a system of coupled differential equations and the between-host dynamics with the classical compartmental models of mathematical epidemiology. Therefore, these models are generally deterministic and do not explicitly incorporate bottleneck size. Recent studies suggest that transmission involves both stochastic and fitness bottlenecks and that some pathogens can begin infection with only a small number of cells in the initial inoculum (Schmid-Hempel and Frank, 2007; Joseph et al., 2015; Moxon and Kussell, 2017). Also, repeated artificial bottlenecks in viral populations have been demonstrated to severely restrict viral fitness (Duarte et al., 1992). This, along with new DNA sequencing techniques, has led to a resurgence of interest in understanding the size and nature of transmission bottlenecks.

The bottleneck size determines how much of the initial diversity from one host passes to another during transmission. Small bottlenecks limit diversity of the founding population in the new host and alter the mutational structure of the population in the recipient, which then can be significantly different than the original host. On the other hand, if the bottleneck is large, transmission does not significantly impact the composition of the founding population so the recipient more closely matches the host (McCrone and Lauring, 2018).

Most experimental studies suggest tight bottlenecks and small founding populations are common, but these results vary significantly depending on the virus, host, and mode of transmission. Experimental infections with tagged influenza clones in ferrets and guinea pigs indicate that airborne transmission imposes a much tighter bottleneck between hosts than by direct contact (Varble et al., 2014). In many cases (Edwards et al., 2006; Keele et al., 2008) HIV has been shown to have a very sharp bottleneck of one or very few cells, consisting of a single genotype. Similarly, HCV has been shown to have bottlenecks of up to only two virions. However, several studies suggest that there are important exceptions to this (Abrahams et al., 2009), which include co-infection by other pathogens that can relax these bottlenecks (Sagar et al., 2004; Haaland et al., 2009). Vertical transmission of HCV between mother and child can significantly increase the bottleneck to between 100 and 184 virions (Fauteux-Daniel et al., 2017). Alternatively, estimates of influenza bottlenecks between humans are much larger and depend on the strain of the virus. A recent experimental study estimates that the mean bottleneck for H1N1 is between 90 and 192 virions and between 114 and 248 virions for H1N3 (Poon et al., 2016). Our work indicates that disease subject to smaller transmission bottlenecks may be subject to stronger forces of selection at the population level than currently thought and this may be a driver of rudimentary types of division of labor (Black et al., 2020).

Our focus in this work has been on simplicity; creating a minimal model to investigate the role of the bottleneck size, but as such the current model has several limitations. The dispersal process is not mechanistic and fully synchronous, which leads to discrete generations. Also, the resource patches are fully isolated so that no migration takes place hence when cells are dispersed to a new patch they all have the same parent patch. In a real system, all of these conditions are likely to be violated to some extent. Dispersal events would likely be initiated by a combination of ecological and biological conditions and take place in a less synchronous environment (Rainey and Kerr, 2010). Many of these complications are implicitly related to spatial structure of the population which plays a role in how dispersal mixes cells between patches. Preliminary investigations show that spatial structure and non-synchronous dispersal can be incorporated into these models, but more work is required to fully understand the implications for the evolutionary dynamics.

Adding a program of growth and development with specific cell types has the potential to align more closely with ongoing experimental studies (Rainey and Kerr, 2010; Rose et al., 2020). A developmental program requires the coordination between cells and could be an important driver of bottleneck size evolution and is something to be explored in further studies. There are also certain aspects of our results that can be traced to the particular model we have implemented rather than being fully general. For example, cells in a patch do not start dying until waste products increase in concentration, so early extinction of cells does not occur. If cells were allowed to die from the beginning of colonisation then many patches would go extinct by chance, especially with smaller bottlenecks and lower growth rates. This may then favour larger bottlenecks which would have some buffer against this. Such effects would require a more ecologically complex model, as stated above.

We have shown that ecological scaffolding can provide a robust framework for evolutionary change even when the size of transmission bottleneck between patches is relaxed. By quantitatively measuring the effects of a larger bottleneck on the forces of selection, a plausible way to further investigate the effects of bottleneck size on evolutionary dynamics in structured populations has been revealed. This may help inform further investigations into viral transmission in addition to providing insights into evolutionary transitions in individuality.

## 7 Acknowledgements

AJB was supported by an Australian Research Council’s DECRA fellowship (DE160100690), which also funded CN’s PhD stipend. PB’s research was supported under the Australian Research Council’s Discovery Projects funding scheme (Project Number DE210100303). PBR thanks the Max Planck Society for generous core support and funding from the Deutsche Forschungsgemeinschaft (DFG) Collaborative Research Center 1182 ‘Origin and Function of Metaorganisms’ (grant no. SFB1182).

