## Supplementary Material for "The Effect of Bottleneck Size on Evolution in Nested Darwinian Populations"

#### 1 Further model details

In this section we give more technical details relating to the the folded state-space modifications and simulation algorithm.

In the original formulation of the model the position within the state vector, indexed by $i$, aligns with the growth rate vector $\boldsymbol{\beta}$, which is populated with all the $n$ values tracked by the model as defined by

$$\beta_i = \beta_1 + \mu(i-1) \quad i \geq 1\,. \tag{1}$$

Figure S1 shows an illustration of the relation between types in this setup.

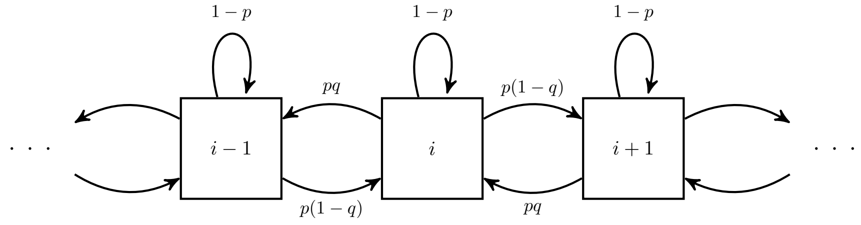


Figure S1: Illustration of the the ordering of types in the first version of the model and how different events create different types. In this version, types are ordered linearly according to their growth rates. Each cell type $i$ replicates with probability $1-p$ (represented by self-loops), can mutate to a type with faster growth rate $\beta_{i+1}$ with probability $p(1-q)$ (represented by arrows to the right) or a type with a slower growth rate $\beta_{i-1}$ with probability $pq$ (represented by arrows to the left).

The folded version orders the states according to the relation

$$\lambda_\ell = \begin{cases} \beta_k - \frac{\mu}{2}(\ell - 1), & \text{if } \ell \text{ is odd} \\ \beta_k + \frac{\mu}{2}\ell, & \text{if } \ell \text{ is even} \end{cases} \qquad \ell \geq 1\,,$$

where $\beta_k$ is chosen dynamically as described in the main text. The ordering of this folded version is illustrated in Figure S2.

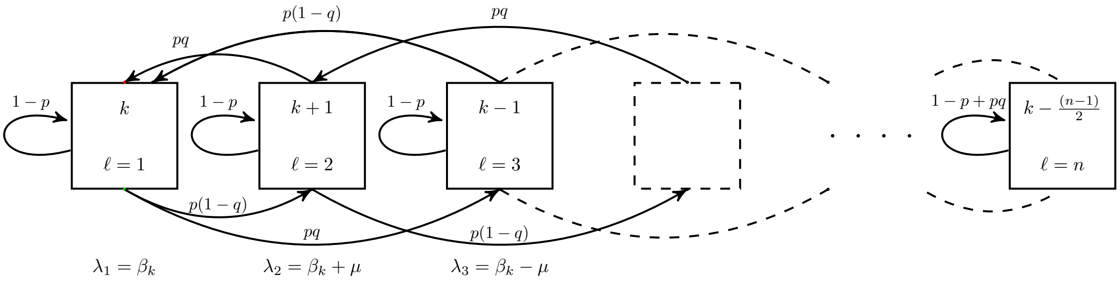


Figure S2: Illustration of the relation between types with the folded ordering of the state vector. The arrows indicate the same set of birth, death and mutation events as in Figure S1.

To further illustrate this construction and the relationship to the original state vector, consider an example model that tracks a total of $n = 9$ different types with $\boldsymbol{\beta} = (2.7, 2.8, \ldots, 3.5)$. Suppose the bottleneck for a patch consists of three distinct types, $\bar{\beta} = \{3.1, 3.0, 2.9\}$. Under the original construction the initial condition would be

$$\mathbf{X}(0) = \Big(0, V, \Big\langle 0, 0, 1, 1, 1, 0, 0, 0, 0 \Big\rangle\Big)\,.$$

However using the folded version, the state vector is centred around the type $\beta_5 = 3.0$. Therefore, the initial condition would be

$$\mathbf{X}(0) = \Big(0, V, \Big\langle 1, 1, 1, 0, 0, 0, 0, 0, 0 \Big\rangle\Big)\,,$$

with

$$\boldsymbol{\lambda} = (3.0, 3.1, 2.9, 3.2, 2.8, 3.3, 2.7, 3.4, 2.6)\,.$$

Figure S3 shows some populations from example simulations under the two different representations.

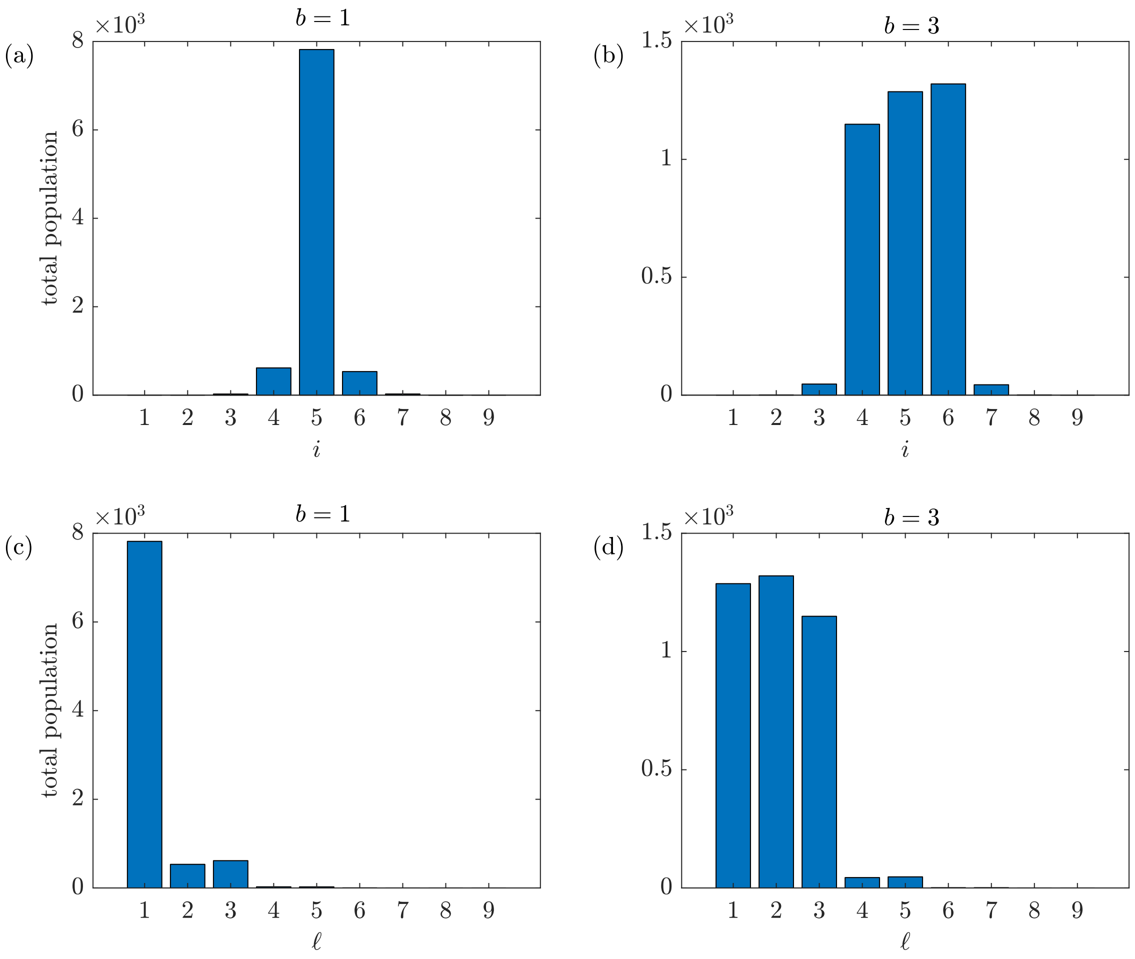


Figure S3: The distribution of types within a patch at dispersal, $T$. The top row is a simulation of the model with state space ordering defined by Equation 1. Results are shown for a bottleneck of $b = 1$ and $b = 3$ cells. The bottom row shows the same results but using the folded ordering. Other parameters: $p = 0.01$, $\mu = 0.1$, $q = 0.5$.

To simulate the within-patch model, we can calculate the propensities of each event more efficiently by defining some matrices that simplify the computation. For each cell type $i$, there are four possible events as shown in Table 1 of the main text. Each event either creates or destroys an $i$-type and we can therefore write the dynamics in terms of these creation / destruction events. Growth rates are incremented by a fixed mutation step size, so there is only mutation from types with a growth rate directly above or below $i$. This structure remains in the reordered state-space defined above and allows computation of the propensities as follows.

First we define the matrix of fixed probabilities, $\mathbf{\Pi}$:

$$\begin{aligned}\mathbf{\Pi} = &\operatorname{diag}_0((1-p), \dots, (1-p)) + \operatorname{diag}_{-1}(p(1-q), 0, \dots, 0) \\ &+ \operatorname{diag}_{-2}(pq, p(1-q), \dots, pq, p(1-q)) \\ &+ \operatorname{diag}_{+1}(pq, 0, \dots, 0) + \operatorname{diag}_{+2}(p(1-q), pq, \dots, p(1-q), pq)\,,\end{aligned}$$

where $\operatorname{diag}_n$ is a diagonal matrix with $n = 0$ the main diagonal, $n > 0$ rows above and $n < 0$ rows below the main diagonal. The last two rows follow from truncation of the state space at $n$ total types:

$$\mathbf{\Pi}_{n-1,j} = (0, \ldots, 0, p - pq, 0, 1 - p + pq, 0)\,, \quad j = 1, \ldots, n\,,$$
$$\mathbf{\Pi}_{n,j} = (0, \ldots, 0, pq, 0, 1 - pq)\,, \quad j = 1, \ldots, n\,.$$

Therefore, the vector used to determine the propensities of each birth or death event is a concatenation of the following two vectors:

$$\boldsymbol{\alpha}_b = \mathbf{\Pi} \cdot \operatorname{diag}\Big(\boldsymbol{\lambda}\Big) \cdot \mathbf{X} \cdot EV^{-1}\,,$$
$$\boldsymbol{\alpha}_d = \operatorname{diag}\Big(\boldsymbol{\gamma}\Big) \cdot \mathbf{X} \cdot DV^{-1}\,,$$

where $\boldsymbol{\gamma}$ is the vector of death rates of cells in a patch.

### 2 Further single-cell bottleneck results

In this section we give some further results on simulations of the evolutionary process assuming a single cell bottleneck and show how the evolutionary dynamics are altered by the various secondary parameters of the model, $p$, $q$, $\mu$, and $M$.

The effect of $p$ on the within-patch dynamics of the model is illustrated in Figure S4. When the mutation probability is small, very few mutants grow within each patch, which results in homogeneous patches composed of the same population from which they were seeded. Larger values of $p$ result in more mutants, both faster and slower growing, that in turn creates more variability in both the patch composition and the size. The smaller the value of $p$ the smaller the overall number of types that have to be tracked within a patch, which increases computational efficiency, but the overall speed of the evolutionary dynamics is then also slowed as there is less variation, requiring simulations to be run over a larger number of generations before equilibrium is reached. This can be seen in Figure S5 that shows the average evolution of the growth rate for different values of $p$. Throughout the paper we choose $p = 0.01$ as reasonable trade off between all these factors.

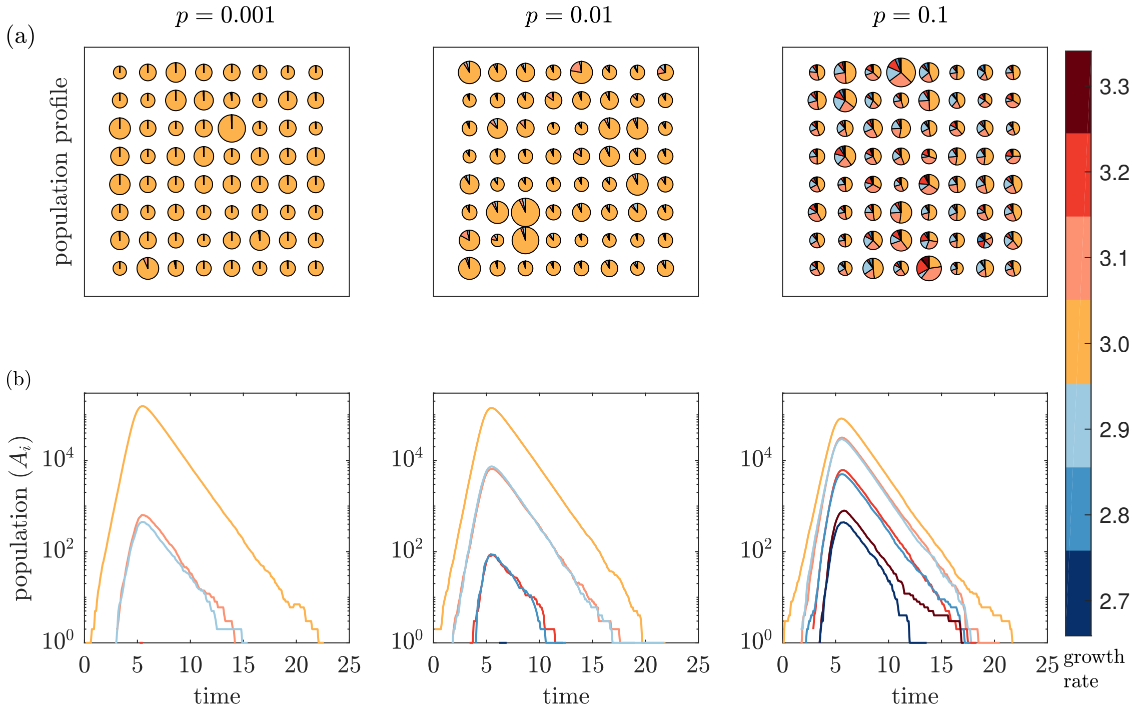


Figure S4: Accumulation of cells at dispersal for different mutation probabilities, $p$. (a) A sample of 64 realisations of the within-patch model each with the same initial condition, $\bar{\beta} = \{3.0\}$, at dispersal time $T = 10$. Each pie has area proportional to the total number of cells with arcs proportional to the composition. (b) A single realisation showing populations by type as a function of time over the growth phase. Both the pie charts and lines are coloured according to the growth rates for each cell type. Other parameters: $q = 0.5$, $\mu = 0.1$.

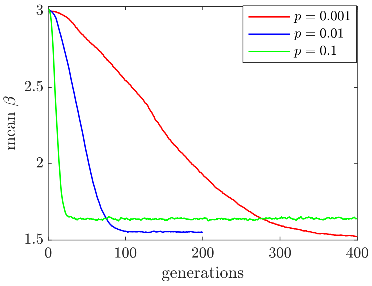


Figure S5: The effects of different values of $p$ on the evolutionary dynamics. Simulations are initialised with a single cell bottleneck, $M = 100$, $\mu = 0.1$ and $q = 0.5$.

The effect of changing the mutation step size, $\mu$, and mutation symmetry, $q$, on the average evolutionary dynamics of the system is shown in Figure S6. Larger $\mu$ implies that the difference in growth rates between types is larger, hence faster growing mutants have a larger advantage within the patch. Therefore, larger $\mu$ results in a faster convergence to the equilibrium growth rate. Similarly, the parameter $q$ controls the symmetry in the production of mutant types. A smaller value of $q$ biases a mutation toward faster-growing mutants, which in turn increases competition within the patch and hence slows the rate of evolution.

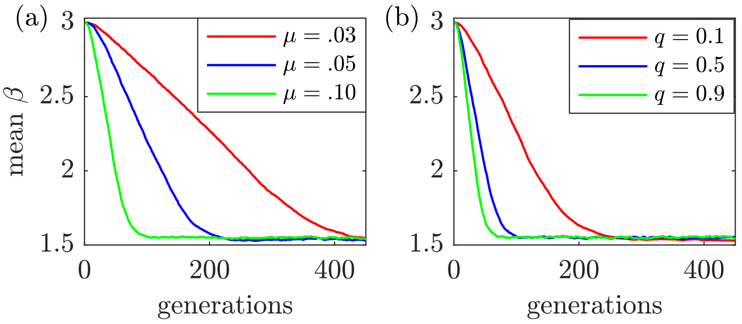


Figure S6: Change in evolutionary dynamics with different values of $\mu$ (a) and $q$ (b). Unless otherwise labelled, parameters are: $\mu = 0.1$, $q = 0.5$, $p = 0.01$.

Figure S7 shows realisations of the model for four different values of $M$, the total patch population. We observe that as the number of patches decreases, individual realisations become noisier and the mean rate of change in the growth rate slows, but the same equilibrium is eventually reached. As $M \to \infty$, the initial average rate of change in the mean growth rate over all $M$ patches over the first 100 generations, $\Delta E[\beta]/\Delta G$, tends to a limiting value. This implies that the inherent randomness in the dispersal and growth processes have a large impact on the evolutionary dynamics, which gets averaged out as $M$ increases. More precisely, the impact of stochasticity on the evolutionary trajectories is reduced when a larger number of patches, $M$, is considered.

Figure S7(b) shows the expected changes in the growth rate due to within- and between-patch forces of selection. Notice that the between-patch selection grows stronger as $M \to \infty$, while within-patch selection remains constant.

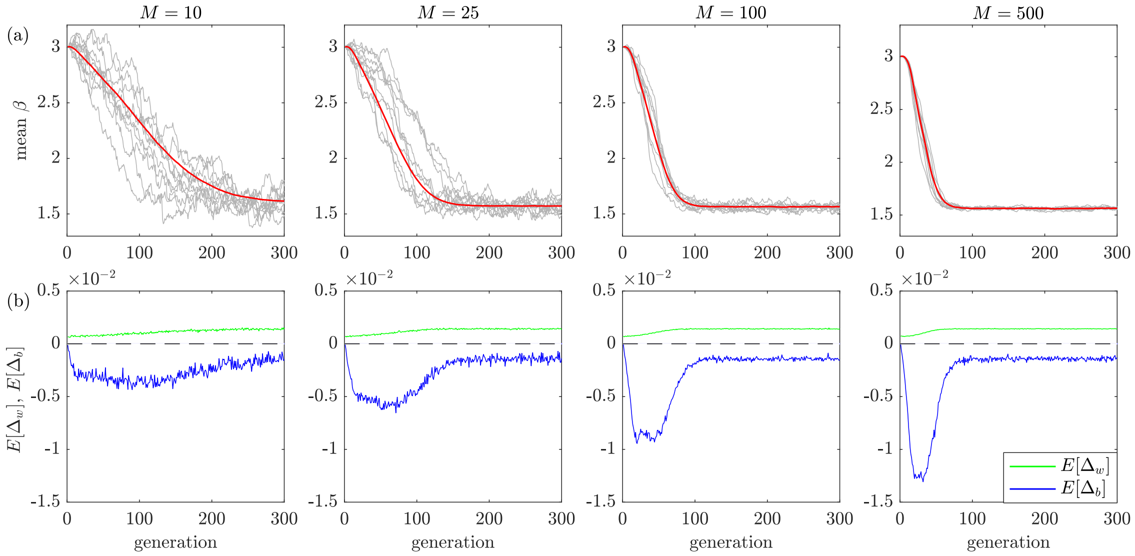


Figure S7: The effect of $M$, the total number of patches, on the evolutionary dynamics. (a) The change in growth rate over a population of patches over generations. The grey lines show several individual realisations and the red lines show the averages over these realisations. (b) The expected change in the growth rate per cell, $\Delta_b$ and $\Delta_w$, as a function of the generation. Each quantity is averaged over a number of realisations. Other parameters: $b = 1$, $\mu = 0.1$, $p = 0.01$, $q = 0.5$

### 3 Further multi-cell bottleneck results

In this section we present some further multi-cell bottleneck results.

One complicating factor to the dynamics described in the main text is that patches that start with more cells (from a bigger bottleneck) will on average have a maximum population that peaks earlier. This phenomenon is illustrated in Figure S8, which shows the mean total population as a function of time for various increasing bottleneck sizes. If populations that originate from larger bottlenecks peak earlier, then at longer times the populations are smaller, hence the optimal growth rate will also be smaller to compensate.

A patch that is colonised by more cells initially will typically enter the exponential growth phase more rapidly, decreasing the stochasiticity in the final patch size at dispersal. This effect is illustrated in Figure S9 which shows the patch size distribution at dispersal for increasing bottleneck sizes. As $b$ is increased the mean population shifts to smaller values, reflecting the dynamics described in the previous paragraph and the variance also decreases. Note that in both Figures S8 and S9 the patches are started from homogeneous populations of cells. In the full simulations with dispersal, colonising cells are not homogeneous, the implications of which are discussed in the main text.

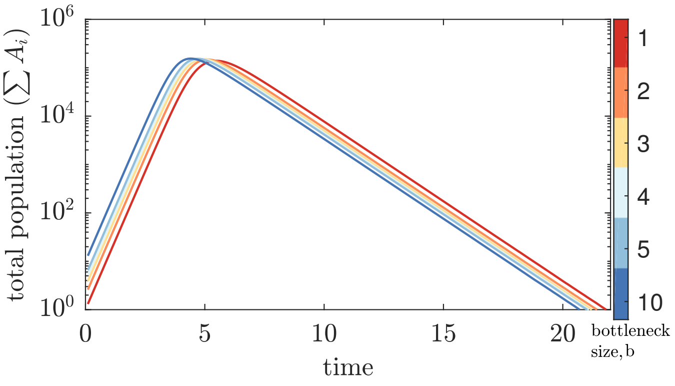


Figure S8: Mean population as a function of time for a range of bottleneck sizes. Each simulation is started from a homogeneous population of cells with growth rate $\beta = 3.0$. As the size of the bottleneck increases, the time for the population to peak is earlier on average. Other parameters: $p = 0.01$, $q = 0.5$ and $\mu = 0.1$.

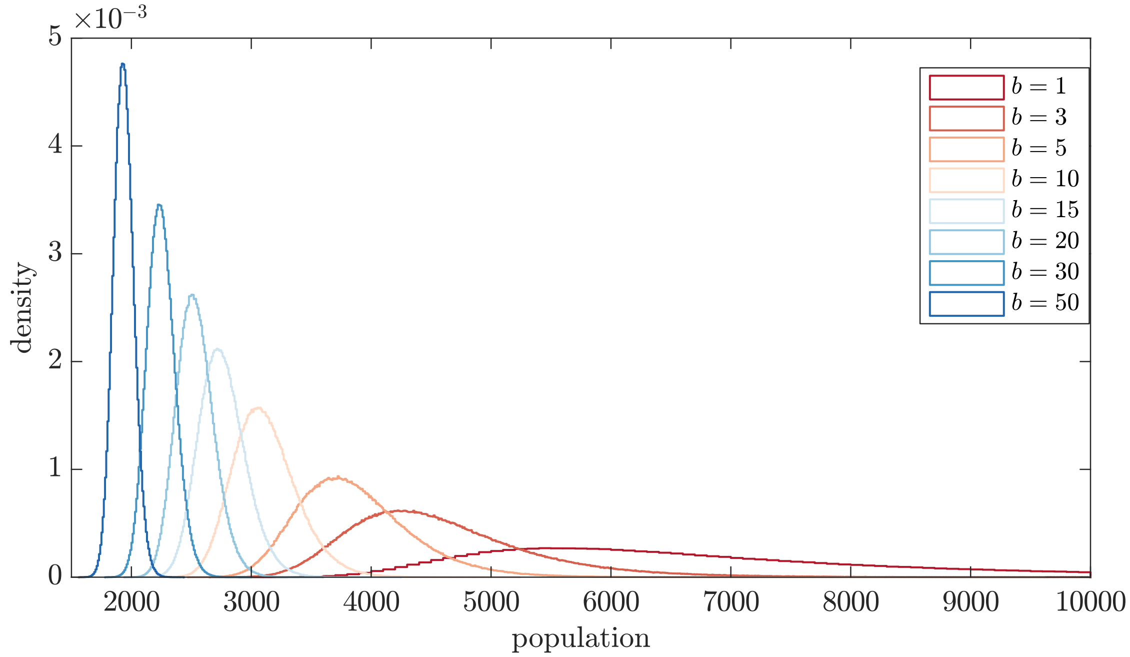


Figure S9: Patch size distributions at a dispersal time of $T = 10$ for increasing bottleneck sizes. As the size of the bottleneck is increased, both the mean and variance of the distribution decrease. Other parameters: $\beta = 3.0$, $p = 0.01$, $q = 0.5$ and $\mu = 0.05$.
